# Semantic Disease Gene Embeddings (SmuDGE): phenotype-based disease gene prioritization without phenotypes

**DOI:** 10.1101/311449

**Authors:** Mona Alshahrani, Robert Hoehndorf

## Abstract

**Motivation:** In the past years, several methods have been developed to incorporate information about phenotypes into computational disease gene prioritization methods. These methods commonly compute the similarity between a disease’s (or patient’s) phenotypes and a database of gene-to-phenotype associations to find the phenotypically most similar match. A key limitation of these methods is their reliance on knowledge about phenotypes associated with particular genes which is highly incomplete in humans as well as in many model organisms such as the mouse.

**Results:** We developed SmuDGE, a method that uses feature learning to generate vector-based representations of phenotypes associated with an entity. SmuDGE can be used as a trainable semantic similarity measure to compare two sets of phenotypes (such as between a disease and gene, or a disease and patient). More importantly, SmuDGE can generate phenotype representations for entities that are only indirectly associated with phenotypes through an interaction network; for this purpose, SmuDGE exploits background knowledge in interaction networks comprising of multiple types of interactions. We demonstrate that SmuDGE can match or outperform semantic similarity in phenotype-based disease gene prioritization, and furthermore significantly extends the coverage of phenotype-based methods to all genes in a connected interaction network.

**Availability:** https://github.com/bio-ontology-research-group/SmuDGE

## 1 Introduction

There is now a large number of available methods for the prioritization or prediction of gene-disease associations (Wang *et al.*, 2011; Zhou and Skolnick, 2016; Natarajan and Dhillon, 2014). Computational methods that predict gene-disease associations use a large number of different features and approaches.

Several approaches to the computational prediction of gene-disease associations are based on the guilt-by-association principle (Gillis and Pavlidis, 2012). Using the guilt-by-association approach relies on prior knowledge of a set of genes associated with a disease *D* and a relatedness measure that compares genes with the set of genes associated with *D*; if a gene is strongly related with respect to the relatedness measure it is suggested as a novel candidate gene. Several measures are used to determine relatedness between genes, with the most prominent ones relying on network associations (Aerts *et al.*, 2006; Lee *et al.*, 2011; Köhler *et al.*, 2008) or some form of functional or phenotypic similarity (Schlicker and Albrecht, 2009). However, as guilt-by-association relies on prior knowledge of disease-associated genes, they can not easily be applied to monogenic diseases, and their applications are, in general, limited to few diseases.

Phenotype-based approaches have been particularly successful in finding candidate genes for Mendelian diseases (Hoehndorf *et al.*, 2011). Phenotype-based approaches compare disease phenotypes to a database of genotype–phenotype associations and suggest candidate genes based on measures of phenotype similarity (Köhler *et al.*, 2009; Hoehndorf *et al.*, 2011; Eilbeck *et al.*, 2017).

The main limitation of phenotype-based approaches, however, is the limited amount of phenotype annotations that are associated with particular genotypes in public databases. In the past, one approach to address this limitation is the use of phenotype associations resulting from animal model experiments and the use of ontologies that can combine phenotypes across species so that animal model and human phenotypes can be compared (Hoehndorf *et al.*, 2011; Chen *et al.*, 2012). While the use of model organisms significantly extends the scope of phenotype-based disease–gene prioritization methods, there is nevertheless only a limited amount of phenotype associations available. In particular, genes for which there are no orthologs in other organisms cannot benefit from cross-species phenotype-based approaches. One possible way in which this challenge could be overcome is to predict phenotypes associated with genes that have no such associations in databases, for example through the use of background knowledge in the form of interaction networks.

We developed SmuDGE, a method that combines phenotype similarity and network similarity to predict gene–disease associations, generates features encoding for phenotype-associations for any gene connected in an interaction network. To achieve this goal, SmuDGE combines phenotype similarity with network-based representation learning and propagates information about phenotype-associations through interaction network connections. We demonstrate that SmuDGE can be used to identify candidate genes of disease through the use of phenotype similarity even if no phenotypes are associated with a gene. SmuDGE is freely available from https://github.com/bio-ontology-research-group/SMUDGE.

## 2 Methods

### 2.1 Data sources and versions

We use the PhenomeNET ontology (Rodriguez-Garcia *et al.*, 2017), downloaded on 10 Jan 2017 from the AberOWL repository (Hoehndorf *et al.*, 2015), as our phenotype ontology because it integrates human and model organism phenotypes and allows them to be compared. We use a dataset of diseases with their phenotypes from the HPO database (Köhler *et al.*, 2014), downloaded on 15 Jan 2017.

Furthermore, we use gene-to-phenotype associations observed in mutant mouse models, downloaded from the Mouse Genome Informatics (MGI) database (Blake *et al.*, 2014) on 12 Jan 2017, and gene-to-phenotype associations derived from gene–disease associations and provided by the HPO database, downloaded on 7 Nov 2017. We further used the interactions provided by STRING (Szklarczyk *et al.*, 2011) version 10. STRING contains both direct and indirect interactions.

Our dataset consists of 7,064 diseases with 78,599 associations to 6,597 distinct phenotypes; 3,526humangenes with 153,575 associationsto 6,058 distinct phenotypes?and 11,696 mouse genes with 200,170 associations to 8,603 distinct phenotypes.

We use human-mouse orthology obtained from MGI on 12 Jan 2017 to identify the human orthologs of mouse genes, and associate mouse gene’s phenotypes with their human orthologs, resulting in 144,360 associations between 9,131 human genes and 8,534 distinct phenotypes.

Furthermore, we map all proteins in the STRING interaction network to their gene identifiers using the mappings provided by STRING. The resulting interaction network between genes consists of 493,041 interactions between 14,753 genes.

### 2.2 Construction of the heterogeneous graphs

We have built two kinds of heterogeneous knowledge graphs to study gene–disease associations. The first knowledge graph utilizes the cross-species PhenomeNET ontology (Rodriguez-Garcia *et al.*, 2017) and characterizes phenotypes of human diseases and mouse models. We associate the human orthologs of the mouse genes with mouse phenotypes, resulting in 144,360 associations between human genes and mouse phenotypes. Furthermore, we construct a second version of that graph in which we use human proteins and assign them with their phenotypes obtained from the HPO database.

The second graph aims to exploit a protein-protein interaction network to generate vector representations for genes which don’t have phenotypes. It consists of the same information as the first type of graph plus the STRING interaction network (Szklarczyk *et al.*, 2011).

### 2.3 Similarity computation and evaluation

We use cosine similarity between two vectors *υ*_1_ and *υ*_2_ to determine the similarity of embeddings:

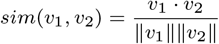

We use cosine similarity to compute the similarity between disease and gene embeddings. We use their similarity as predictor for genes’ having an association with a disease.

As baseline for comparison, we use a semantic similarity measure which exploits the background knowledge in an ontology. We use Resnik’s semantic similarity measure (Resnik *et al.*, 1999) with the Best Match Average (BMA) strategy for combining similarities between individual classes. Resnik’s semantic similarity measure is defined as:

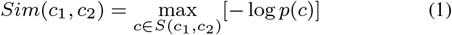

where *c*_1_ and *c*_2_ are the two classes between which similarity is computed, and *S*(*c*_1_, *c*_2_) is the set of superclasses of both *c*_1_ and *c*_2_ in the ontology hierarchy.

To evaluate the performance of the similarity-based predictions, we compute a similarity matrix which contains the pairwise similarities of genes and diseases. For each disease, we rank genes in descending order of the similarity score. We then evaluate at which rank we identify a gene–disease association in our evaluation dataset. As this method results in a ranking classifier (as genes are ranked for each disease), we quantify the performance of the predictions through the area under the receiver operating characteristic (ROC) curve (Fawcett, 2006). A ROC curve is a plot of the true positive rate (TPR) as a function of the false positive rate (FPR). The TPR at a particular rank is defined as a rate of correctly predicted gene-disease associations at this rank, and the FPR is the rate of predicted associations that are not gene–disease associations. As we do not have true negative gene–disease associations, we treat unknown gene–disease associations as negatives.

### 2.4 Supervised prediction and evaluation

SmuDGE is an unsupervised method to generate feature vectors for genes and diseases based on their phenotypes. Using these features in a supervised manner can improve the prediction of associations between two vectors. For this reason, we use an artificial neural network (ANN) to train pairs of gene-disease associations. In this experiment, we use the known disease-gene association as the positive set, and randomly select an equal number of the non-associated disease-gene pairs as the negative set.

For the the training and testing, we perform 5-fold cross validation. We generate folds by sampling diseases, not gene–disease pairs. 80% of the diseases are used for training the ANN and we use the 20% of disease feature vectors for testing. As positive pairs, we then combine the diseases in each fold with the genes associated with the diseases. The aim of this sampling strategy is to guarantee that the ANN does not learn to recognize gene-disease associations for a disease *D* based on genes known to be associated with *D*, and therefore determine how well our method predicts genes associated with diseases if no prior knowledge is available.

We used 10% of training set as a validation set to guide and stop the training if the loss increases in the validation set; alternatively, training will stop after 100 epochs. We use a Rectified Linear Unit as an activation function for the hidden layers (Nair and Hinton, 2010) and a sigmoid function as the activation function for the output layer; we use cross entropy as loss function in training, and Rmsprop (Root Mean Square Propagation) (Hinton *et al.*, 2012) to optimize the neural networks parameters in training.

For the evaluation of the ROCAUC, we create an embedding matrix for each disease in which we fix the first part of the matrix to represent a particular disease embeddings and the second part represents the all gene embeddings. We then apply the learned model on the matrix and rank the genes based on the probability scores. The TPR and FPR at each rank are used to identify the proportion of correctly and falsely predicted associations at each rank respectively.

## 3 Results

### 3.1 Heterogeneous representation of genes, diseases, and phenotypes

In our method we use a knowledge graph as data structure in which we represent genes, diseases, and the phenotypes with which they are associated. Genes, diseases, and phenotypes are represented as nodes in the graph. Edges between phenotypes represent axioms in the Web Ontology Language (OWL) (Grau *et al.*, 2008; Rodriguez-Garcia and Hoehndorf, 2018). We represent diseases using their identifiers from the Online Mendelian Inheritance in Men (OMIM) (Amberger *et al.*, 2011)) database, human genes using their Entrez gene identifier, and phenotypes using the cross-species phenotype ontology PhenomeNET (Rodriguez-Garcia *et al.*, 2017). We connect diseases and genes to the phenotypes they are associated with using the *has phenotype* relation. Additionally, we represent interactions between genes and their products using an *interacts with* relation. We consider all *interacts with* edges as symmetric (i.e., it x interacts-with y then y interacts-with x) and all other edges as non-symmetric.

We associate all OMIM diseases with their phenotypes from the Human Phenotype Ontology (HPO) database (Köhler *et al.*, 2017), and obtain information about interactions between human genes from the STRING database (Szklarczyk *et al.*, 2011).

We build two knowledge graphs which differ in the associations between genes and their phenotypes. In the first case, we use the phenotypes associated with human genes in the HPO database; these phenotypes are indirectly derived from gene–disease associations and the disease phenotypes, i.e., if a gene *G* is associated with a disease *D* and the disease has a set of phenotypes *P*, then all phenotypes in *P* are assigned to *G*. Because this assignment of phenotypes to human genes can indirectly encode for gene–disease associations, we also use mouse model phenotypes as an independent dataset of phenotypes. For this purpose, we identify the phenotypes of non-conditional loss of function mutations (i.e., knockouts) of gene *G* in the Mouse Genome Informatics (MGI) database (Blake *et al.*, 2014); if we can identify a human ortholog of *G*, we assign all phenotypes of *G* to the human ortholog. Phenotypes of mouse models are encoded using the Mammalian Phenotype Ontology (MP) (Smith *et al.*, 2004); through the use of the cross-species PhenomeNET ontology (Rodriguez-Garcia *et al.*, 2017), the phenotypes encoded using MP (in the mouse) and HPO (in human disease) can be compared directly.

Our graphs consist of 7,066 nodes for diseases, 15,391 nodes for human genes, and 9,293 nodes for mouse models phenotypes, 7,833 nodes for human phenotypes. It has 78,599 disease–phenotype associations. Using phenotypes from HPO, we include 153,575 gene–phenotype associations; using mouse phenotypes from MGI, we include 144,360 gene–phenotype associations. We also include 493,041 interactions between genes, all of which we consider as symmetric. Figure 1 illustrates the knowledge graph we generate.

**Fig. 1.**
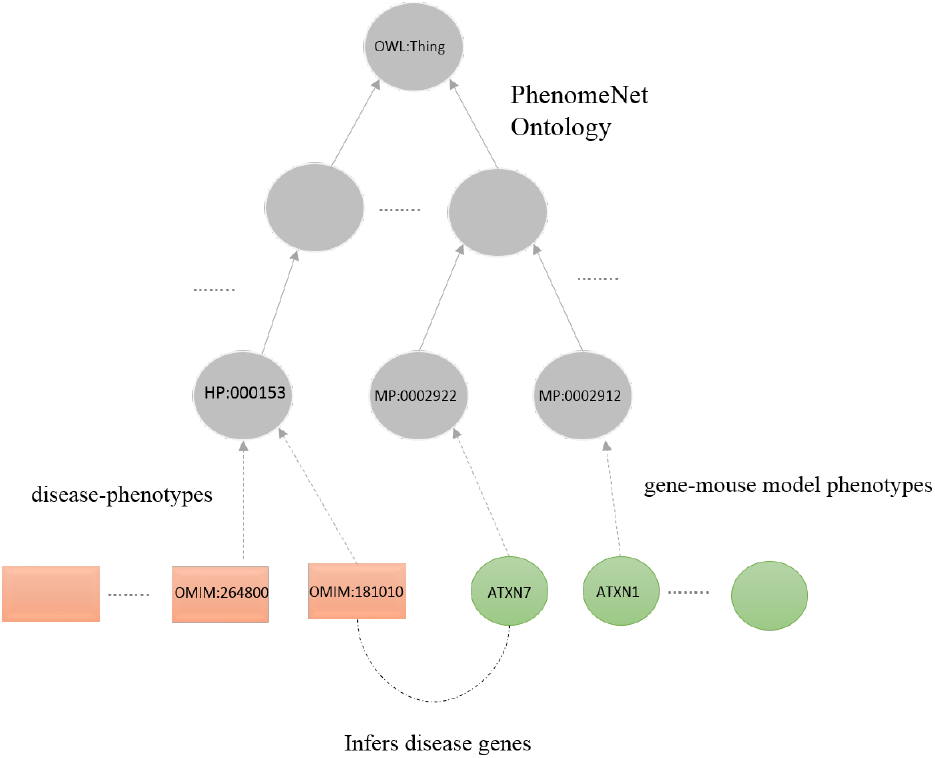
Our knowledge graph consists of gene–phenotype associations (encoded using either HPO or MP, disease–phenotype associations (encoded using the HPO), and the PhenomeNET ontology.

### 3.2 Joint representation learning from PPI network structure and phenotype annotations

We designed an algorithm to encode features based on the phenotypes that are associated with entities in the knowledge, either diseases or genes and gene products, in the form of a dense vector; the vector representation of the genes can then be used in unsupervised or supervised machine learning approaches or other predictive models.

Our algorithm, Semantic Disease Gene Embeddings (SmuDGE), comes in two forms. First, it encodes for the phenotypes that are directly associated with an entity (i.e., a disease or gene/gene product); for this purpose, it generates a dense representation of an entities ontology–based annotations and its superclasses. This algorithm is applicable to all diseases and genes that are directly associated with phenotypes. However, while diseases are commonly associated with (or even defined by) a set of phenotypes, the majority of genes are not associated with phenotypes, neither in humans, where phenotypes are generally derived from gene–disease associations (Köhler *et al.*, 2014), nor in the mouse where phenotypes are the result of phenotyping experiments (de Angelis *et al.*, 2015). Therefore, we use a second form of our algorithm which is applicable to genes without any phenotype associations and in which a phenotype-based representation is assigned indirectly using the network environment in which a gene product is embedded.

For the first version of our algorithm, we ignore all interactions between genes and their products and focus only on encoding an entity’s phenotype annotations and the ontology structure (i.e., subclass relations). Given an entity (disease or gene) *E* that has *has phenotype* edges to the phenotypes *P*_1_,…, *P_n_*, we generate n sentences starting with *E*. We generate *n* sentences where each sentence starts with *E* followed by *P_i_* and all of the superclasses of *P_i_*, for all *P_i_* that are directly associated with *E*.

We then apply a Word2Vec skipgram model (Mikolov *et al.*, 2013) to learn vector representations for each token occurring in a generated sentence, in particular for all entities and phenotype classes. The vectors generated for entities through this approach encode for directly associated phenotypes and all their superclasses, and we call the vectors *P-Vecs* (for Phenotype Vectors).

We can only generate P-Vecs for genes and gene products with directly associated phenotypes. For all other genes, however, we can use their interaction network environment to assign phenotypes that are overrepresented in the neighboring nodes. For a gene *G*, we use a random walk, starting at *G*, over the network of interactions between genes and gene products to randomly sample *G*’s network neighborhood. We terminate the walk once we found a node *G*’ with *has phenotype* edges, or after a pre-determined step limit, whichever occurs first. If the step limit has been reached, we restart the walk at *G*. If the walk found a *G*^’^ with an outgoing *has phenotype* edge, and the phenotypes associated with *G*’ are *P*_1_,…, *P_m_*, then we randomly sample one phenotype *P* of *P_i_*,…, *P_m_* and generate a sentence starting with *G* followed by *P* and all superclasses of *P*; after adding the sentence to our corpus, we restart at *G*. The aim of this approach is to sample the network environment in which *G* is located for phenotypes. Through inclusion of the ontology hierarchy in the generated sentences, the approach is intended to be more robust to differences in specific phenotypes. Similarly to generating P-Vecs, we apply a Word2Vec skipgram model on the generated sentences to produce vector representations of all entities and phenotypes in the corpus. Because these representations are generated from a gene node’s network environment, we call the vectors *E-Vecs* (for Environment Vectors). Figure 2 provides a high-level overview over our method.

**Fig. 2.**
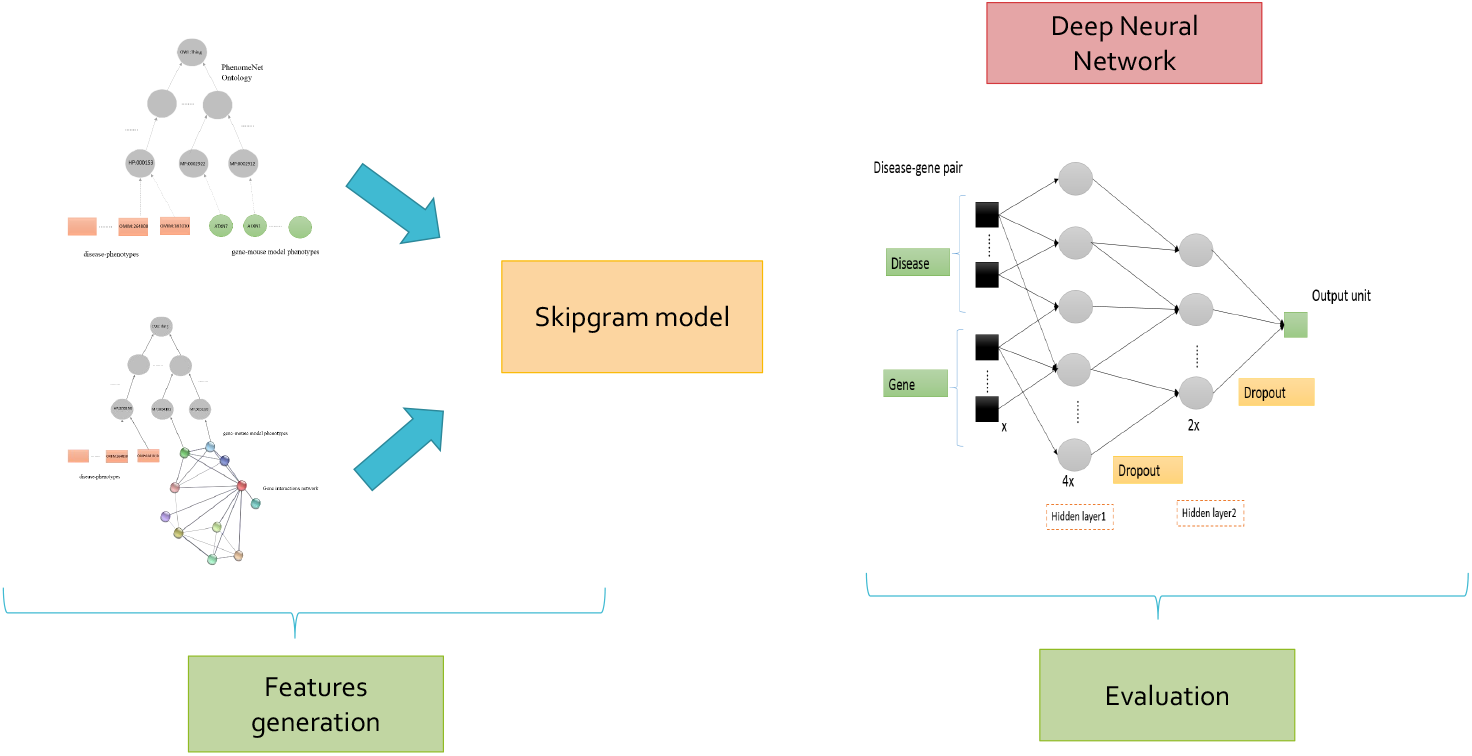
On the left side we show the graphs of disease–phenotype associations, gene–phenotype associations and the PhenomeNET ontology (top), and the graph used to generate E-Vecs at the bottom. We then use a skipgram model to generate vectors for genes and gene products in the graph. These vectors can then be used as input to a similarity measure, or a neural network, to predict interactions or other types of biological relations.

We generate both P-Vecs and E-Vecs for our two graphs (using human and mouse gene–phenotype associations separately). We generate P-Vecs for all human diseases, and we further generate P-Vecs both for human and mouse genes that have phenotypes associated. We generate E-Vecs both for nodes which do not have phenotypes associated and for nodes which have phenotypes associated; if a gene node has directly associated phenotypes, we mask them during the generation of sentences (i.e., random walks) for the E-Vec approach so that only the network environment is sampled for phenotypes. Furthermore, we generate vector representations for all phenotype classes from HPO and MP which are either used to directly annotate a gene or diseases, or are a superclass of a direct annotation.

In total, we generate 9,293 vectors for phenotype classes from MP, 7,833 for phenotype classes from HPO, 7,064 for diseases from OMIM, 9.131 P-Vecs for genes using mouse phenotypes (i.e., assigning phenotypes of mouse genes to their human orthologs), and 3,526 P-Vecs for genes using human phenotype data (i.e., genes-phenotypes associations provided by the HPO database). We generate E-Vecs for 14,753 genes, i.e., for all genes in our interaction network that are connected to a gene with phenotype associations.

### 3.3 Similarity-based prediction of disease-associated genes

We use the generated vectors representing genes and diseases to predict gene–disease associations based on phenotype similarity. The SmuDGE vectors encode phenotype annotations connectivity patterns along with the ontology super classes associated with each phenotype annotation. Similar phenotypes for both genes and diseases indicate similar features vectors and therefore we can infer disease-gene associations by comparing feature vectors.

To determine similarity, we use the cosine similarity measure, and we compute the pairwise similarity between all diseases and genes (using either their P-Vec or E-Vec representation). We then rank the most similar genes for each disease and determine how well we recover known gene–disease associations from OMIM using a receiver operating characteristic (ROC) curve (Fawcett, 2006); we quantify the performance of the similarity-based prediction using the area under the ROC curve (ROCAUC) which is equivalent to the probability that a randomly chosen positive sample is ranked higher than a randomly chosen negative sample. We limit our evaluation of P-Vec similarity to the genes for which we can generate the representations, i.e., 3,526 genes using human phenotypes and 9.131 genes using mouse phenotypes. For comparison, we use a semantic similarity measure to compare disease and gene phenotype annotations. For the evaluation of E-Vec similarity, since we generate representations for all of the genes in the interactions network, we evaluate the disease vectors against the set of 14,753 genes vectors. Figure 3 shows the results using P-Vec similarity for human and mouse phenotypes, and Table 1 shows the results using E-Vec similarity based on human and mouse phenotypes.

**Fig. 3.**
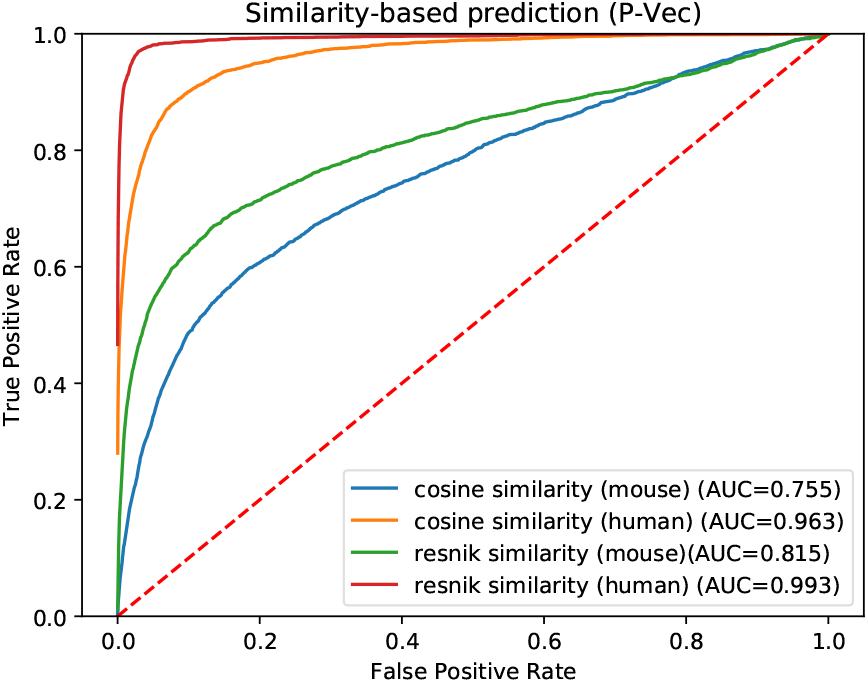
ROCAUC for gene–disease association using cosine similarity (P-Vec)

**Table 1.** ROCAUC results for E-Vecs and P-Vecs using cosine similarity and ANN.

| Evaluations | P-Vec | E-Vec |
| --- | --- | --- |
| Human phenotype (cosine) | 0.96 | 0.77 |
| Mouse phenotype (cosine) | 0.75 | 0.69 |
| Human phenotype (ANN) | 0.97 | 0.77 |
| Mouse phenotypes (ANN) | 0.84 | 0.81 |

We find that P-Vec similarity using human phenotypes results in almost perfect prediction (ROCAUC 0.96), which is the consequence of how the phenotypes have been assigned to genes in the HPO database (i.e., the phenotypes are identical to the phenotypes of the disease with which the gene is associated, and using them for prediction is therefore almost circular); these similarities are therefore not truly predictive but mainly reproduce our evaluation dataset. However, using mouse phenotypes, we obtain a ROCAUC of 0.75 when comparing P-Vecs to the disease vectors. Using E-Vecs, we obtain a ROCAUC of 0.77 when using human gene–phenotype associations and a ROCAUC of 0.69 using gene–phenotype associations from the mouse. Notably, because we mask all direct gene–phenotype associations when generating E-Vecs, our use of human gene–phenotype associations does not encode for gene-disease associations, and the performance of this similarity-based evaluation is therefore indicative of predicting disease-associated genes in the absence of gene–phenotype associations.

### 3.4 Supervised prediction of disease-associated genes

Cosine similarity can only be applied to vectors of the same dimension, and furthermore cannot easily account for dataset-specific features. Therefore, we also apply supervised machine learning to “learn” a function (akin to a similarity measure) that takes two phenotype representation vectors as input and is predictive of gene–disease associations. We use an artificial neural network (ANNs) to learn these functions in a supervised manner; the ANN layout we use is shown in Figure 4.

**Fig. 4.**
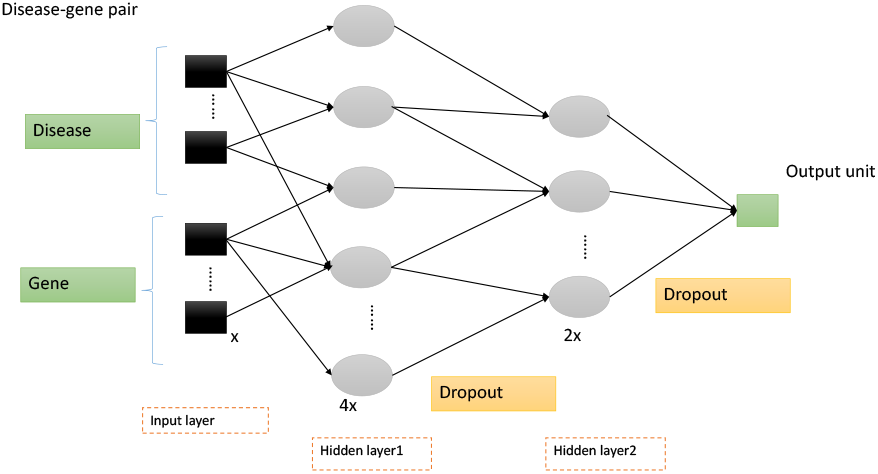
The ANN model we built, the input is the pair of disease-gene feature vector of size x, the first hidden layer consists of 4x hidden units, and the second hidden layer consist of 2x hidden units, we use a dropout of 0.5 to mitigate the effects of overfitting. We optimize the number of hidden layers and units, the dropout ratio, the input embeddings dimensions. For clarity in the figure, some ANN connections are omitted.

Several approaches to computational prediction of gene-disease associations utilize the principle known as “guilt-by association” (Gillis and Pavlidis, 2012) which infers the associations of a gene to a disease based on the similarity to other genes associated with the disease. As a result, it fails to predict genes for diseases with no prior knowledge of any associated genes. Supervised training to predict gene-disease associations is similar to the guilt-by-association approach if some genes associated with a disease have been used in training and the model is evaluated on the remaining genes, because knowledge about disease-associated genes is used to predict more associations. To estimate the performance of our method for predicting gene-associations for diseases without associated genes, we first split the training and testing based on the diseases, not on gene-disease pairs. In particular, we select 80% of the diseases and all their associated genes for training, and apply the model to predict all the genes for the remaining 20% of the diseases.

We evaluate each type of vector representation individually using our ANN classifier approach (see Materials & Methods). As in the similarity–based prediction of disease-associated genes, we use pairs of P-Vecs or E-Vecs as input to the ANN and use the ANN to compute the “similarity” between them as a predictor of a gene–disease associations. The ROC curves for using mouse vectors, as well as the comparison to a semantic similarity measure, are shown in Figure 5 and Table 1. We find that using the ANN significantly improves the results compared to the unsupervised, similarity-based approach, increasing the ROCAUC from 0.75 to 0.84 for mouse phenotypes and P-Vecs as well as using E-Vecs from 0.69 to 0.81 ROCAUC.

**Fig. 5.**
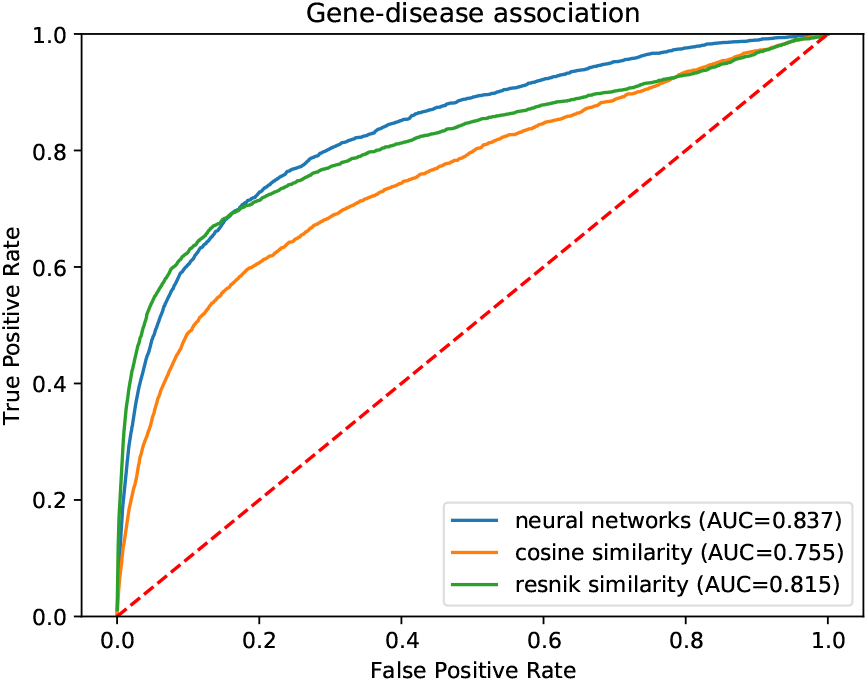
Comparision of ROCAUC for gene-disease association using a neural network, cosine similarity and Resnik’s semantic similarity measure using mouse phenotypes and SmuDGE’s P-Vec approach for generating feature vectors.

As another use case, we also evaluated how well SmuDGE can predict gene-disease associations for diseases with only a single association compared to diseases with multiple associated genes. Although our dataset is stratified by disease and known associations are therefore not used during training of the neural network, we intend to test the performance of our approach on rare diseases for which no or only little information is available. Figure 6 and Figure 7 show the result. We find that for diseases which have more gene associations and are likely better studied, SmuDGE can predict associated genes better than for diseases with only a single associated gene, using both the P-Vec and E-Vec approach.

**Fig. 6.**
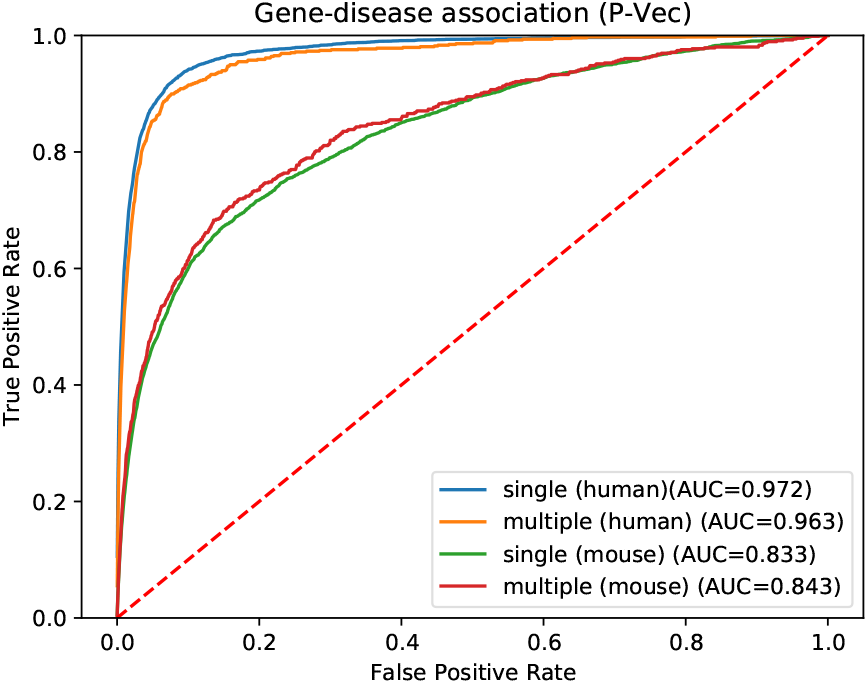
ROCAUC for predicting gene–disease associations for genes with a single or multiple associated genes using SmuDGE’s P-Vec approach.

**Fig. 7.**
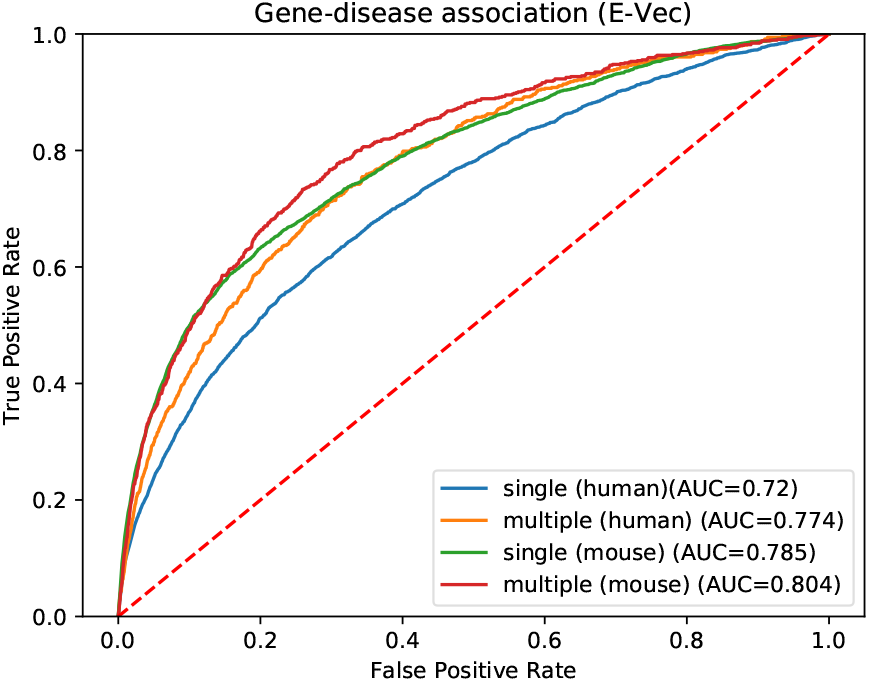
ROCAUC for predicting gene–disease associations for genes with a single or multiple associated genes using SmuDGE’s E-Vec approach.

## 4 Discussion

SmuDGE is an algorithm that exploits ontologies and knowledge graphs to learn representations of genes, gene products, and diseases, based on the phenotypes they are associated with. While we demonstrate in our evaluation that the performance of SmuDGE in predicting gene-disease associations matches, and sometimes outperforms, traditional phenotypes–based gene prioritization methods such as PhenomeNET (Hoehndorf *et al.*, 2011) or the MouseFinder (Chen *et al.*, 2012), we see our main contribution in extending the phenotype- and similarity-based approaches for gene–disease prioritization to all genes represented in an interaction network (or knowledge graph).

The prediction of disease genes using phenotype-similarity has been highly successful (Köhler *et al.*, 2009; Hoehndorf *et al.*, 2011; Chen *et al.*, 2012) and a major limitation has been the availability of phenotypes for many genes. The use of non-human model organisms such as the mouse (Meehan *et al.*, 2017) can generate phenotype representations of human genes even in the absence of clinically determined phenotypes associated with a gene; however, even using model organism phenotypes, phenotype similarity can still not be applied to a large portion of human genes due to missing data or lack of a non-human ortholog.

SmuDGE’s E-Vecs don’t use directly associated phenotypes but encode for phenotypes of a gene based on knowledge of interactions – both direct and indirect – of a gene with other genes with which phenotypes are associated. The disease vector representations we generate always encode for phenotypes, and the success in identifying gene–disease associations when comparing both demonstrates that E-Vecs and disease phenotype vectors have similar (and possibly complementary) information, sufficient for their comparison to be predictive of disease mechanisms (i.e., disease-associated genes).

The E-Vecs we construct in our work further encode, although indirectly, for phenotypic network modules, since they are generated through random walks on an interaction network and will encode for phenotypes overrepresented within a network region. In future work, we plan to evaluate whether our approach can identify interacting genes that may be jointly associated with a disease, such as in digenic and other oligogenic diseases (Gazzo *et al.*, 2016). We may also explore the possibility to combine SmuDGE with variant prioritization tools to provide additional information that can be used to determine whether a variant is associated with a phenotype or not (Boudellioua *et al.*, 2017).

## Funding

This work was supported by funding from King Abdullah University of Science and Technology (KAUST) Office of Sponsored Research (OSR) under Award No. URF/1/3454-01-01 and FCC/1/1976-08-01.

